# The human Ska complex and Ndc80 complex interact to form a load-bearing assembly that strengthens kinetochore-microtubule attachments

**DOI:** 10.1101/245167

**Authors:** Luke A. Helgeson, Alex Zelter, Michael Riffle, Michael J. MacCoss, Charles L. Asbury, Trisha N. Davis

**Affiliations:** Department of Biochemistry, University of Washington, Seattle, WA 98195; Department of Genome Sciences, University of Washington, Seattle, WA 98195; Department of Physiology and Biophysics, University of Washington, Seattle, WA 98195

## Abstract

Accurate segregation of chromosomes relies on the force-bearing capabilities of the kinetochore to robustly attach chromosomes to dynamic microtubule tips. The human Ska complex and Ndc80 complex are outer-kinetochore components that bind microtubules and are required to fully stabilize kinetochore-microtubule attachments *in vivo*. While purified Ska complex tracks with disassembling microtubule tips, it remains unclear whether the Ska complex-microtubule interaction is sufficiently strong to make a significant contribution to kinetochore-microtubule coupling. Alternatively, Ska complex might affect kinetochore coupling indirectly, through recruitment of phospho-regulatory factors. Using optical tweezers, we show that the Ska complex itself bears load on microtubule tips, strengthens Ndc80 complex-based tip attachments, and increases the switching dynamics of the attached microtubule tips. Crosslinking mass spectrometry suggests the Ska complex directly binds Ndc80 complex through interactions between the Ska3 unstructured C-terminal region and the coiled-coil regions of each Ndc80 complex subunit. Deletion of the Ska complex microtubule-binding domain or the Ska3 C-terminus prevents Ska complex from strengthening Ndc80 complex-based attachments. Together our results indicate that the Ska complex can directly strengthen the kinetochore microtubule interface and regulate microtubule tip dynamics by forming an additional connection between the Ndc80 complex and the microtubule.

**SIGNIFICANCE STATEMENT:** Microtubules are dynamic, tube-like structures that drive the segregation of duplicated chromosomes during cell division. The Ska complex is part of a molecular machine that forms force-bearing connections between chromosomes and microtubule ends. Depletion of the Ska complex destabilizes these connections and disrupts cell division. The Ska complex binds microtubules but it is unknown if it directly holds force at microtubules or indirectly stabilizes the connections. Here, we show that the Ska complex makes a direct force-bearing linkage with microtubule ends and assembles with another microtubule binding component, the Ndc80 complex, to strengthen its ability to withstand force. Our results suggest that the Ska and Ndc80 complexes work together to maintain the connections between chromosomes and microtubule ends.

## INTRODUCTION

Depolymerizing spindle microtubules generate forces required to separate duplicated chromosomes during mitosis. The kinetochore couples dynamic microtubule ends to chromosomes and harnesses the energy released by depolymerizing microtubules to pull duplicated chromosomes to opposite poles. Kinetochore-microtubule attachments must sustain piconewton-scale loads, especially during metaphase when bioriented kinetochores are subject to tension from opposing spindle microtubules. Attachments that are too strong or too weak can generate erroneous chromosome-microtubule attachments and promote chromosome mis-segregation during cell division (1). The incorrect segregation of chromosomes leads to aneuploidy and has been linked to chromosomal instability (2, 3). The attachment strength established and maintained between kinetochores and dynamic microtubule ends is fundamental to faithful chromosome segregation and cell division.

*In vivo* experiments show that the heterotrimeric Ska complex (Ska1, Ska2 and Ska3; Figure 1A) is important for the stability of kinetochore-microtubule coupling and suggest at least three models for how it might contribute to coupling strength. Purified Ska complex binds directly to microtubules *in vitro* (4), and loss of Ska complex *in vivo* delays mitotic progression and has been associated with chromosome congression failure and mitotic cell death (5–8). Based on these observations, one view is that the Ska complex contributes directly to kinetochore-microtubule coupling (6, 8, 9). However, some studies suggest instead that the Ska complex plays a more indirect, regulatory role in kinetochore-microtubule coupling by recruiting protein phosphatase 1 to the kinetochore, rather than by bearing microtubule-generated forces (10). Ska complex localizes to kinetochores *in vivo* through interactions with the Ndc80 complex (Hec1, Nuf2, Spc24 and Spc25; Figure 1A), an essential component of the kinetochore-microtubule interface (11–14). This observation raises a third possibility, that the Ska complex might enhance Ndc80 complex-based coupling independently of its own microtubule binding affinity (15). Purified Ska complex alone tracks with depolymerizing microtubule tips (6) and has also been found to enhance the microtubule lattice binding and tip-tracking of the Ndc80 complex (16). While these findings are consistent with a direct role for Ska complex in kinetochore tip-coupling, they do not address the load-bearing capacity of Ska complex-based attachments. Thus, it remains uncertain whether the Ska complex can bear significant load on microtubule ends, either alone or in combination with the Ndc80 complex.

**Figure 1.**
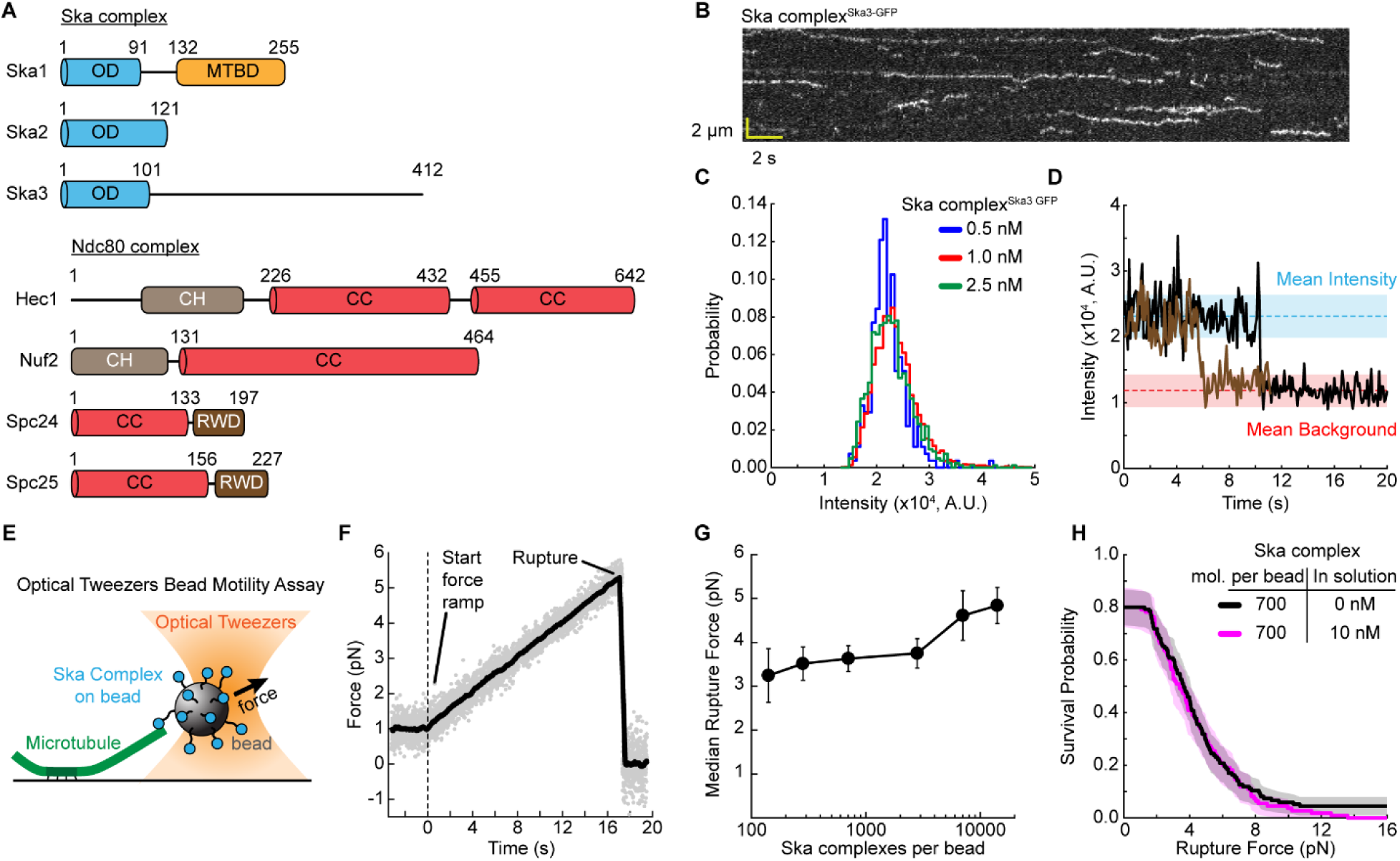
Ska complex bears load on microtubule ends. (A) Domain architecture of the Ska complex and Ndc80 complex. OD: oligomerization domain; MTBD: microtubule binding domain; CH: Calponin homology domain; CC: coiled-coil; RWD: RING finger, WD repeat, DEAD-like helicases domain. (B) Example kymograph of Ska complex^Ska3-GFP^ molecules binding a microtubule. (C) Histogram of tracked Ska complex^Ska3-GFP^ particle intensities for three different concentrations. (D) Two example intensity versus time traces of tracked Ska complex^Ska3-GFP^ particles. After loss of particle tracking, due to dissociation or bleaching, the background was sampled for several frames to calculate the background intensity. Blue dashed line indicates the mean particle intensity for all tracked molecules; red dashed line indicates the mean background intensity. Shaded regions are standard deviation. (E) Cartoon of the optical tweezers based bead motility assay with Ska complex attached to the beads. A bead coated in Ska complexes is bound to the end of a dynamic microtubule. Using the optical tweezers, a force is applied that pulls on the microtubule Ska complex connection. (F) Raw data of a Ska complex rupture force experiment (gray). Black line is data smoothed using a 50-point sliding window. Vertical dashed black line indicates start of force ramp. (G) Median rupture force versus Ska complex on the bead. Error bars are standard deviation from bootstrapping analysis of the median. The median values and errors are calculated from the same data shown in Supplemental Figure S2A. (H) Rupture force survival probability plot for 700 Ska complex molecules per bead without (black) and with (magenta) 10 nM Ska complex in solution. Shaded areas are 95% confidence intervals from Kaplan-Meier analysis.

Here, we tested the microtubule-end, load-bearing strength of the human Ska and Ndc80 complexes, both together and independently. We found that Ska complex bears load at microtubule ends on its own and strengthens Ndc80 complex-based end attachments. Using crosslinking mass-spectrometry, we found that the Ska3 unstructured C-terminal region of Ska complex interacts with the coiled-coil regions of the Ndc80 complex. Furthermore, we show that strengthening Ndc80 complex-based attachments requires the Ska complex to simultaneously bind the Ndc80 complex and the microtubule. Our results suggest the Ska complex and Ndc80 complex directly interact with each other and with microtubules to form a multipartite load-bearing assembly that strengthens kinetochore-microtubule attachments.

## RESULTS

### Ska complex bears load on microtubule ends

The Ska complex is reported to dimerize in solution and to cooperatively bind the microtubule lattice as a dimer or as higher order oligomers (6, 16, 17). Before measuring the strength of its attachments to microtubules, we used TIRF microscopy to examine the oligomeric state of the Ska complex at the low nanomolar concentrations used in microtubule binding and rupture force assays. Individual particles of GFP-tagged Ska complex (Ska complex^Ska3-GFP^) bound and diffused along Taxol-stabilized microtubules, as reported previously, and similarly to the lattice diffusion of other kinetochore components (Figure 1B) (16, 18–20). The mean residence time of Ska complex^Ska3-GFP^ particles on microtubules was 5.2 ± 0.1 s, similar to previously measured residence times (Figure S1A) (16, 18). Particle intensities fell within a unimodal, approximately-gaussian distribution that did not change across a 5-fold increase in concentration, and they photobleached or dissociated in single steps (Figure 1C-D). Moreover, individual Ska complex^Ska3-GFP^ particles, when bound sparsely onto coverslip surfaces, exhibited single-step photobleaching and their mean intensity before bleaching matched that of single GFP-tagged yeast Ndc80 complexes (Figure S1B). Using SEC-MALS, we confirmed that Ska complex^Ska3-GFP^ in solution can form a dimer and exists in a monomer-dimer equilibrium at micromolar concentrations (Figure S1C) (17). However, our TIRF data suggests that at low nanomolar concentrations, the Ska complex binds the microtubule lattice as a single complex.

Using an optical-tweezers bead motility assay, we next measured the microtubule end-binding strength of the Ska complex. We coated beads with the Ska complex at various concentrations, to control the surface density of the molecules on the bead (Figure 1E). Depending on the surface density and molecular structure, one or more molecules can simultaneously interact with the microtubule tip, an arrangement that mimics the multi-valency at kinetochore-microtubule interfaces *in vivo* (21, 22). Individual Ska complex-coated beads were first attached to the growing tips of single microtubules anchored to a coverslip. After an initial low force was applied and the bead was verified to track with tip growth, the force was increased gradually until the attachment ruptured. Median rupture strengths for populations of Ska complex-coated beads were 3 to 5 pN, depending on the surface density (Figure 1F,1G, S2A and Table S1). These observations show that tip-couplers based on purified Ska complex alone can bear significant loads.

Previous work shows that Ndc80 complex microtubule attachments are strengthened through avidity (19). Increasing the surface density on the beads increases the number of Ndc80 complexes that can simultaneously reach the microtubule end (see below and (19)). To test if the Ska complex behaves similarly, we measured the strength of Ska complex-based attachments as a function of its surface density on beads. We observed only a small, 1.5-fold increase in Ska complex attachment strength over a 100-fold range in surface density; whereas the strength of human Ndc80 complex-based attachments increased more substantially, by 4.2-fold over a 24-fold density range (Figure 1G, and see Figure 3E below). Furthermore, addition of 10 nM free Ska complex in solution did not increase attachment strength of bead-bound Ska complex, consistent with the lack of Ska complex oligomerization at nanomolar concentrations (Figure 1H). Taken together, our data show the Ska complex is load-bearing and suggest that its load-bearing capacity is largely established at low molecular surface densities and not strongly enhanced by additional Ska complexes.

### Ska complex Ska3 C-terminus is not required for load-bearing

To identify interacting regions between the Ska complex and microtubules, we performed crosslinking mass spectrometry of Ska complex incubated with Taxol-stabilized microtubules. In agreement with previous reports, we observed crosslinks between microtubules and the Ska1 C-terminal microtubule binding domain (MTBD) as well as between microtubules and the Ska3 unstructured C-terminus (residues 102-402) (Figure 2A, S3) (23, 24). To test the importance of these regions for load-bearing interactions between the Ska complex and microtubules, we measured the attachment strength of mutant Ska complexes missing either the Ska1 MTBD (Ska1 ΔMTBD) or the Ska3 C-terminus (Ska3 ΔC) (Figure 2B). Beads coated with mutant Ska complex^Ska1 ΔMTBD^ failed to bind to microtubules, indicating that the MTBD is required for formation of a load-bearing attachment. In contrast, the fraction of beads coated with Ska complex^Ska3 ΔC^ that bound microtubules was similar to wild-type (Figure 2C) and their end attachment strength was only slightly reduced (by 1.3 fold, Figure 2D and Table S1). These observations confirm that, within the Ska complex, both Ska1 and Ska3 interact with microtubules. The Ska1 MTBD is necessary for load-bearing interactions with microtubules whereas the Ska3 C-terminus makes only a minor contribution.

**Figure 2.**
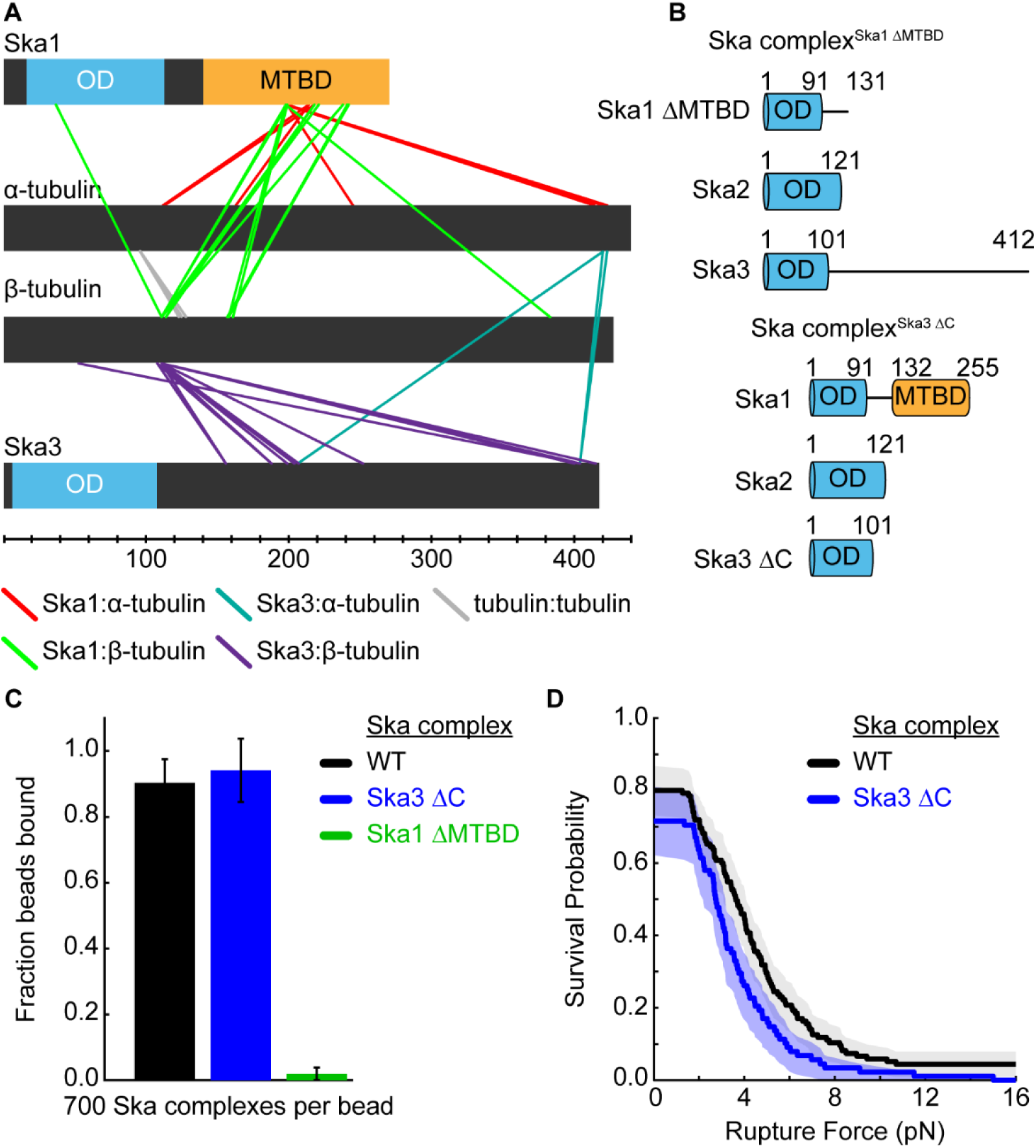
Ska3 C-terminus is not required for Ska complex load-bearing on microtubule ends. (A) Crosslinks identified between tubulin and Ska1 or Ska3. Crosslinking reaction with Ska complex and Taxol-stabilized microtubules was performed for 30 minutes with the amine to carboxyl crosslinker EDC. Intra-Ska complex and Ska2 crosslinks are not shown for clarity; see Figure S3 for all crosslinks identified and http://proxl.yeastrc.org/proxl/viewProject.do?project_id=49 for all data. (B) Domain architecture of the Ska complex mutants. OD: oligomerization domain; MTBD: microtubule binding domain. (C) Fraction of Ska complex coated beads that bound to microtubules: Wild-type Ska complex (black), Ska complex^Ska3ΔC^ (blue), Ska complex^Ska1ΔMTBD^ (green). Error bars are counting uncertainty. (D) Rupture force survival probability plot for 700 Ska complex molecules per bead (black) and 700 Ska complex^Ska3ΔC^ mutant molecules per bead (blue). Shaded areas are 95% confidence intervals from Kaplan-Meier analysis.

### Ska complex strengthens Ndc80 complex-based microtubule attachments

Ska complex increases the affinity of Ndc80 complex for the microtubule lattice and can promote Ndc80 complex tip tracking in the absence of force (16). To determine if the Ska complex can increase the load-bearing capacity of the Ndc80 complex, we measured the rupture force of Ndc80 complex-based attachments with and without the Ska complex added free in solution (Figure 3A). Adding the Ska complex strengthened Ndc80 complex-based microtubule end attachments when the Ndc80 complex was at a low surface density on the beads, but not when it was at a high density (Figure 3B, 3E, S2B and Table S1). The increase in strength afforded by the Ska complex at low Ndc80 complex surface density was greater than the rupture strength of the Ska complex alone, suggesting a synergistic effect. These results show that the Ska complex strengthens Ndc80 complex-based coupling, particularly when the latter is weak due to low avidity.

**Figure 3.**
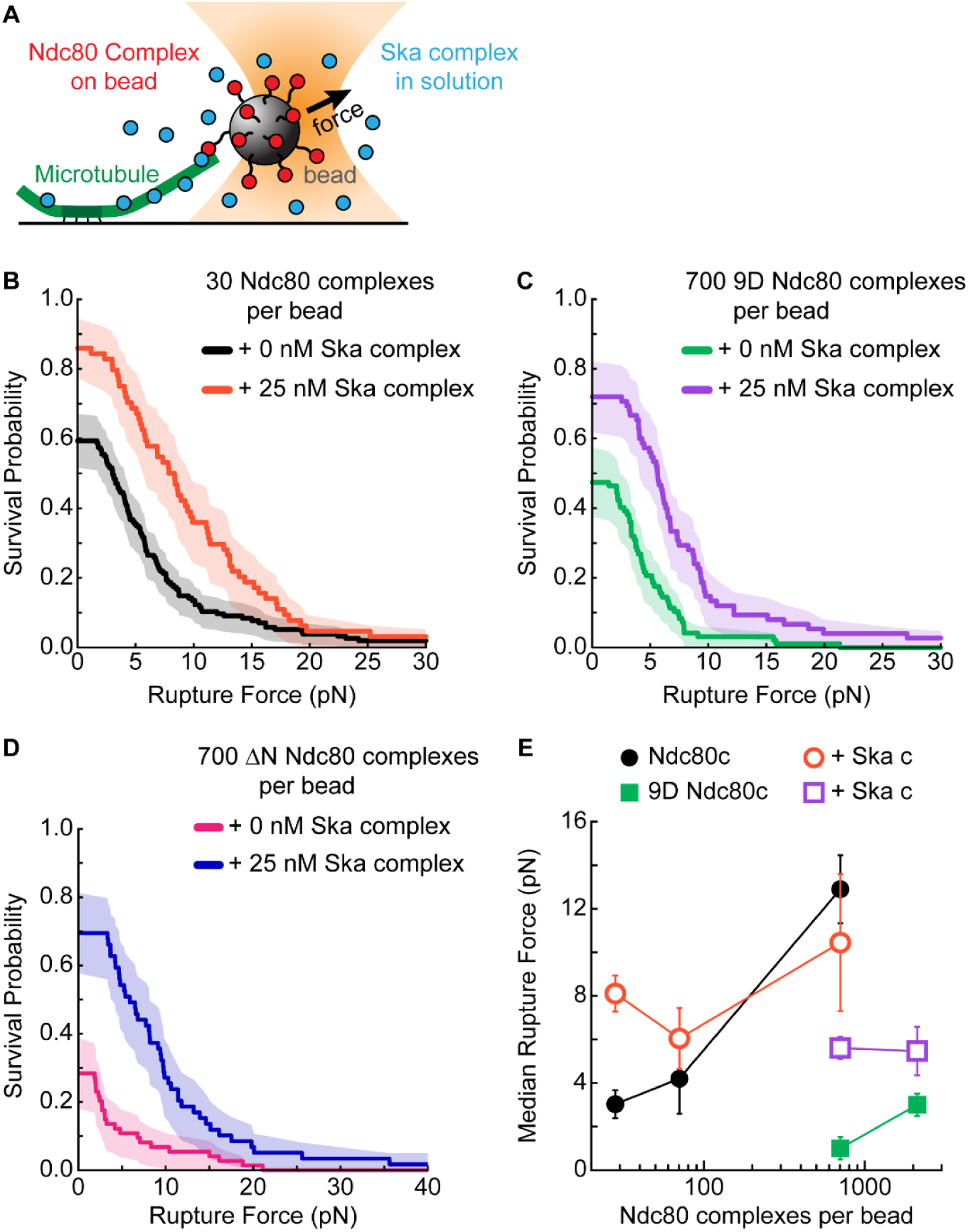
Ska complex strengthens Ndc80 complex microtubule attachments. (A) Schematic of the optical tweezers based bead motility assay with Ndc80 complex attached to the bead and Ska complex free in solution. (B) Rupture force survival probability plot for 30 Ndc80 complex molecules per bead without (black) and with (orange) 25 nM Ska complex in solution. (C) Rupture force survival probability plot for 700 9D Ndc80 complex molecules per bead without (green) and with (purple) 25 nM Ska complex in solution. (D) Rupture force survival probability plot for 700 ΔN Ndc80 complex molecules on the bead without (magenta) or with (blue) 25 nM Ska complex in solution. All shaded regions on survival probability plots are 95% confidence intervals from Kaplan-Meier analysis. (E) Median rupture force versus Ndc80 complex per bead. Error bars are standard deviation from bootstrapping analysis of the median. Closed symbols are Ndc80 complex on the bead and open symbols are Ndc80 complex on the bead with Ska complex in solution. For clarity in panel E, the median data from panel D was omitted from the median versus concentration plot, see Supplemental Table S1 for all median values. In panel E, The median values and errors are calculated from the same data sets shown in B, C and Supplemental Figure S2 B, C.

Next, we tested if the Ska complex could strengthen Ndc80 complex-based attachments that were weakened due to a decreased affinity between the Ndc80 complex and the microtubule. We introduced Aurora B phosphomimetic mutations (serine/threonine to aspartate) in all 9 phosphorylation sites in the Hec1 N-terminal tail to generate the mutant, 9D Ndc80 complex. These mutations dramatically decrease the affinity of the Ndc80 complex for microtubules (20, 25, 26). As expected, we found the mutant 9D Ndc80 complex formed attachments that were significantly weaker than those formed by wild-type Ndc80 complex (Figure 3C). Adding free Ska complex increased the attachment strength of the Ndc80 complex 9D mutant by > 5-fold (Figure 3C, 3E). We raised the surface density of the mutant 9D Ndc80 complex on the beads by 3-fold and found that the Ska complex could also moderately strengthen the attachments formed at this higher density of the 9D Ndc80 complex (Figure 3D, S2C). Furthermore, we tested a mutant Ndc80 complex lacking the entire unstructured N-terminal 80 amino acid tail of Hec1 (ΔΝ Ndc80 complex). As expected, this mutant ΔΝ Ndc80 complex formed weak attachments on its own that, just like the 9D mutant, could be strengthened by the addition of free Ska complex (Figure 3D, 3E). Together, these results show that the Ska complex strengthening is independent of the Hec1 N-terminal tail.

Purified yeast Ndc80 complex and native yeast kinetochore particles detach more frequently from disassembling tips than from assembling tips (19, 27). We verified that this difference also occurs for human Ndc80 complex by applying a force-clamp. Beads coated with human Ndc80 complex were attached to growing tips and then subjected to a constant tension of ~2 pN. Under this condition, the Ndc80 complex-based couplers tracked continuously with end growth and shortening, remaining persistently attached as the tips switched spontaneously between assembling and disassembling states (Figure 4A, 4B). The mean detachment rate for Ndc80 complex-based couplers from disassembling tips was 14-fold higher than from assembling tips, confirming that the coupling was less stable during tip disassembly (Figure 4C, 4D and Table S2). Interestingly, adding Ska complex in solution specifically stabilized the coupling during tip disassembly, reducing the detachment rate 2-fold, with no apparent affect during assembly. Altogether, these results show that Ska complex enhances Ndc80 complex-based attachment in several situations where the coupling would otherwise be relatively poor: when avidity is reduced by lowering the number of participating Ndc80 complexes, when affinity is reduced by adding phosphomimetic mutations in the Hec1 tail or removing the tail, or when attachments are intrinsically destabilized by disassembly of the microtubule tip.

**Figure 4.**
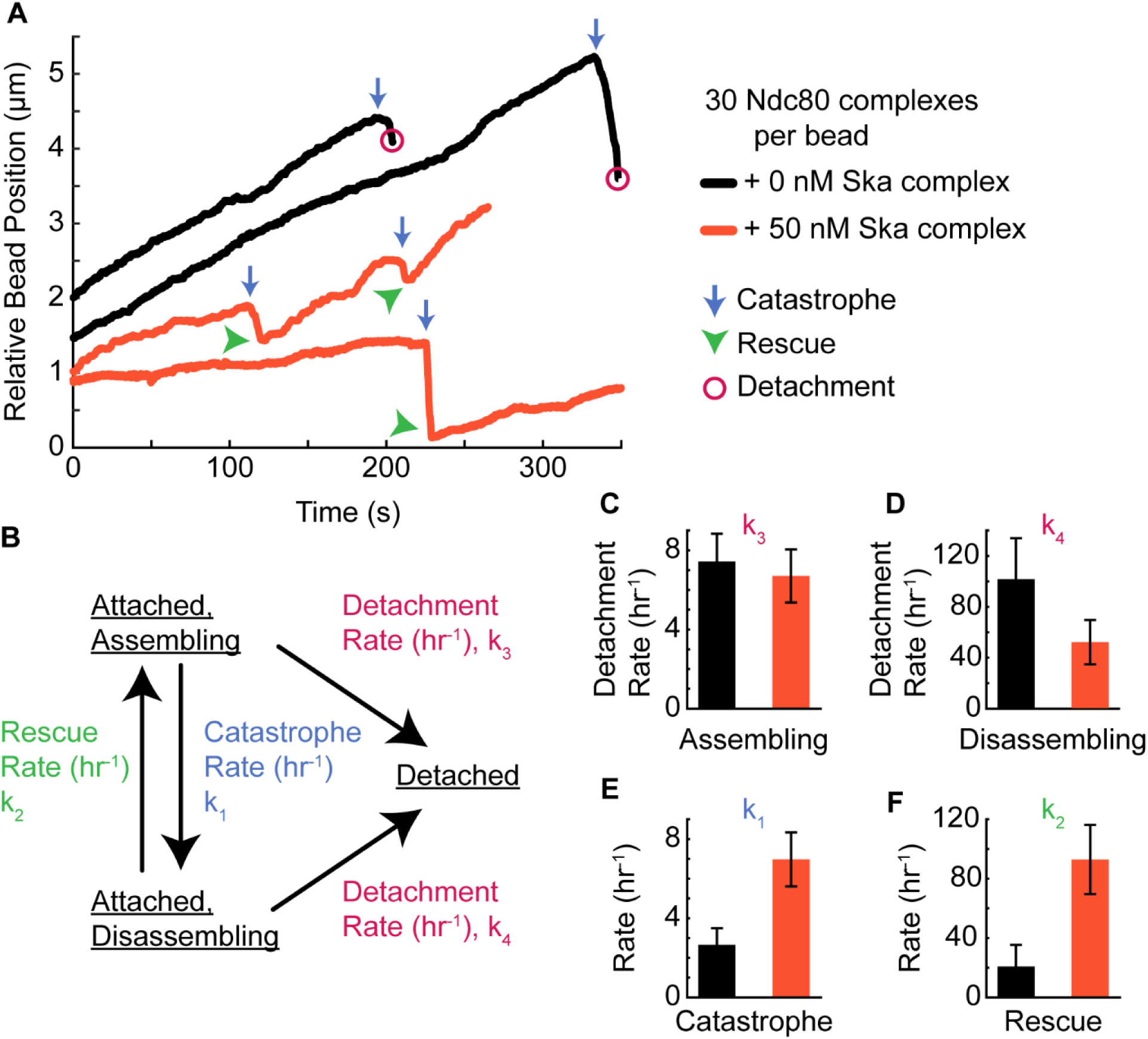
Ska complex affects the dynamics of Ndc80 complex and bound microtubules. (A) Four example bead position versus time traces for 30 Ndc80 complex molecules per bead without (black) and with (orange) 50 nM Ska complex in solution. An average force of 2 pN was exerted on the bead. Blue arrows indicate catastrophe events; green arrowheads indicate rescue events; magenta open circles indicate detachment events. For clarity, the starting position of each trace is offset by an arbitrary amount. (B) Model of coupler microtubule detachment rates and microtubule switching rates measured from constant force bead tracking experiments. (C-F) Measured rates for 30 Ndc80 complex molecules per bead without (black) and with (orange) 50 nM Ska complex in solution. Measured rates are: (C) Detachment rate from an assembling microtubule; (D) Detachment rate from a disassembling microtubule; (E) Catastrophe rate; (F) Rescue Rate. Error bars are counting uncertainty.

### Ska complex changes how the Ndc80 complex governs microtubule switching behavior

Upon alignment at the metaphase plate, chromosomes oscillate between poleward and anti-poleward motions which are partially driven by the switching kinetics of the kinetochore microtubules (28, 29). Altering the microtubule binding affinity of Ska or Ndc80 complexes, independently, dampens these metaphase oscillations *in vivo* (16, 30). To test whether couplers based on the Ndc80 and Ska complexes can affect microtubule tip switching *in vitro*, we measured the dynamics of tips coupled to Ndc80 complex-decorated beads under a constant force, with or without Ska complex in solution (Figure 4A, 4B and Table S2). Indeed, the rescue rate for tips attached to Ndc80 complex-based couplers increased 4.5-fold upon addition of free Ska complex (Figure 4D). This observation is similar to previous findings showing that microtubule rescue rates increase as Ndc80 complex attachments are strengthened (20). Moreover, addition of Ska complex increased the catastrophe rate for attached tips by 2.7-fold (Figure 4C). These results show that the Ska complex changes how the Ndc80 complex governs microtubule behavior and suggests that together they may increase the switching frequency of kinetochore-bound microtubules.

### Ska complex binds the Ndc80 complex coiled-coil through the Ska3 C-terminus

Multiple studies suggest that the Ska complex and Ndc80 complex interact directly, but the interaction interface between the complexes has not been defined (14, 31). To identify the specific regions involved in their interaction, we performed crosslinking mass spectrometry with Ska complex, Ndc80 complex and Taxol-stabilized microtubules. The Ska3 unstructured C-terminus (residues 102-412) crosslinked robustly with the Ndc80 complex and microtubules (Figure 5A, S4). 328 unique crosslinks were found between the Ndc80 and Ska complexes. Of these, 97% (318 of 328) were between Ska3 and the Ndc80 complex, distributed across the Ska3 C-terminus and amongst all four Ndc80 complex subunits. Ska3 primarily crosslinked to regions of the Ndc80 complex that are predicted to form coiled-coils. Few Ska3 crosslinks were observed with the CH domains of Hec1 and Nuf2 or the RWD domains of Spc24 and Spc25. These results suggest that the Ndc80 complex and Ska complex directly interact through the Ska3 unstructured C-terminus that preferentially binds to coiled-coil regions throughout the Ndc80 complex.

**Figure 5.**
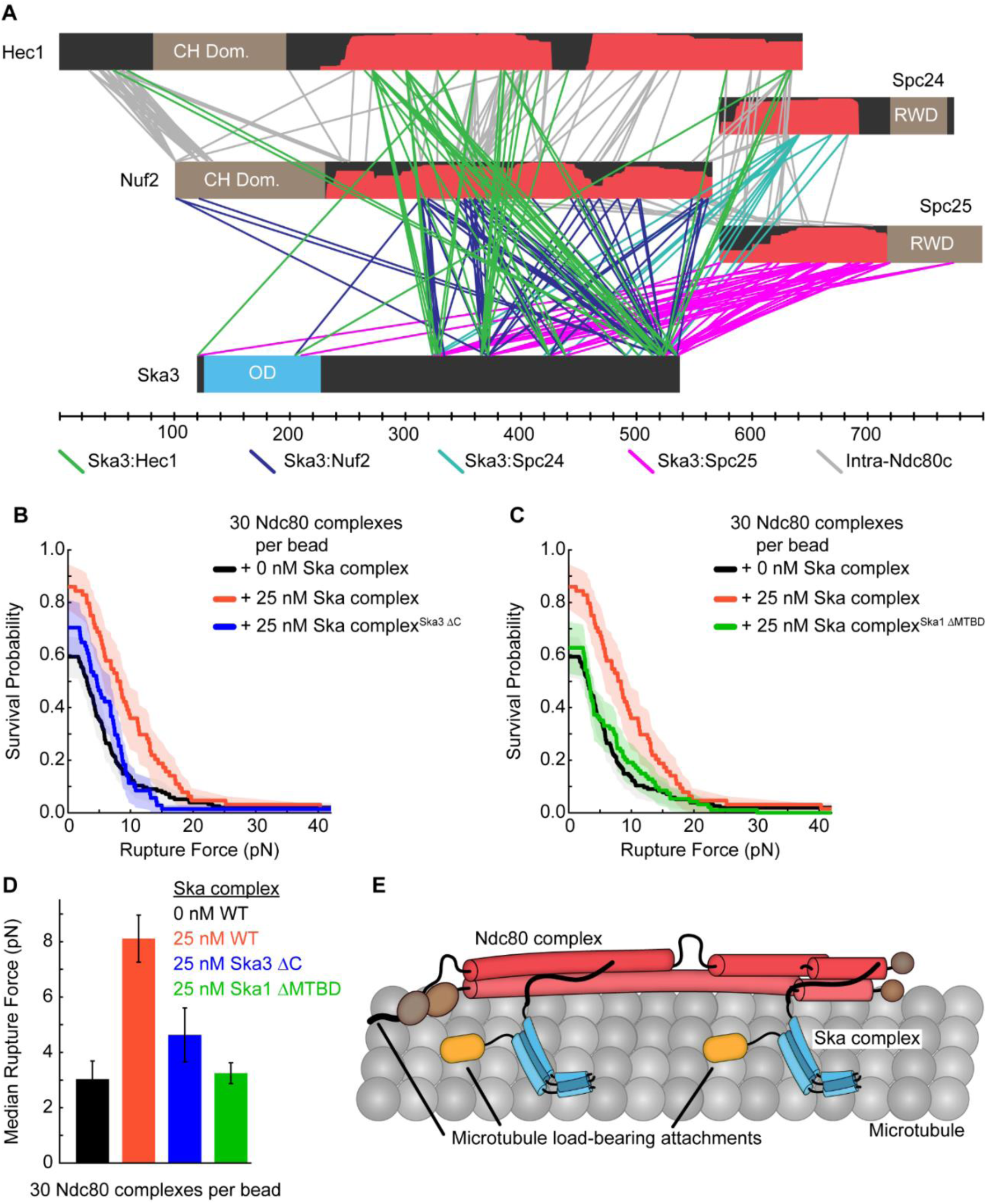
Ska complex must bind Ndc80 complex and microtubules to strengthen attachments. (A) Crosslinks identified between Ska3 and Ndc80 complex. Crosslinking reaction with Ska complex, Ndc80 complex and Taxol-stabilized microtubules was performed for 15 minutes with the amine to amine crosslinker BS3. Ska1, Ska2 and tubulin crosslinks are not shown for clarity; see Figure S4 for all crosslinks identified and http://proxl.yeastrc.org/proxl/viewProject.do?project_id=49 for all data. Red shaded regions indicated predicted coiled-coil (Paircoils2) with probability scores from 0.8-1.0. (B and C)Rupture force survival probability plot for 30 Ndc80 complex molecules per bead without Ska complex (black, data repeated from Figure 3B) and with either: 25 nM Ska complex wild-type (orange, data repeated from Figure 3B) or Ska complex^Ska3 ΔC^ mutant (blue) or Ska complex^Ska1 ΔMTBD^ mutant (green). All shaded regions on survival probability plots are 95% confidence intervals from Kaplan-Meier analysis. (D) Median rupture force for 30 Ndc80 complex molecules per bead with the indicated Ska complexes; colors based on B and C, WT = Wild-type Ska complex. Error bars are standard deviation from bootstrapping analysis of the median. The median values and errors are calculated from the same data sets shown in B and C. (E) Possible model of how Ska complex and Ndc80 complex directly interact to form multiple microtubule load-bearing attachments.

### The Ska complex and Ndc80 complex must bind each other and microtubules to strengthen Ndc80 complex-based attachments

The Ska complex is capable of binding directly to both the Ndc80 complex and to microtubules (6, 13, 14). We have shown that the Ska complex enhances Ndc80 complex-based coupling. Together these observations suggest that Ska complex might form an extra linkage between the Ndc80 complex and the microtubule. However, it is possible that the Ska-dependent enhancement of Ndc80 complex-based coupling occurs indirectly, where Ska complex affects microtubule tip structure in a way that enhances tip-binding of the Ndc80 complex. To test this possibility, we measured the strength of Ndc80 complex-based tip-couplers after addition of a truncated Ska complex, missing the major Ndc80 interaction site within the Ska3 C-terminus (Ska complex^Ska3 ΔC^). Deletion of the Ska3 C-terminus nearly abolished the ability of the Ska complex to strengthen the Ndc80 complex-based tip-attachments (Figure 5B, 5D and Table S1), indicating that direct binding of the Ndc80 and Ska complexes is required for strengthening.

If the Ska complex enhances Ndc80 complex-based coupling by forming an extra linkage between the Ndc80 complex and the microtubule, then removing the major microtubule binding domain of the Ska complex should abolish the enhancement. Indeed, the mutant Ska complex^Ska1 ΔMTBD^ was unable to strengthen Ndc80 complex attachments (Figure 5C, 5D). Crosslinking mass spectrometry with the mutant Ska complex^Ska1 ΔMTBD^ found abundant crosslinks between Ska3 and the Ndc80 complex, similar to wild-type, suggesting that the mutant Ska complex^Ska1 ΔMTBD^ retains normal interactions with the Ndc80 complex (Figure S5). Together, these results support a model where Ska complex strengthens Ndc80 complex-based tip-attachments by binding the Ndc80 complex directly and providing an additional load-bearing bridge to the microtubule (Figure 5E).

## DISCUSSION

Previous studies have established that depletion of the Ska complex *in vivo* generally weakens kinetochore-microtubule attachments, thereby: 1) diminishing the numbers of attachments that are resistant to cold treatment (6, 8, 9), 2) causing more frequent kinetochore detachments during congression (32) and 3) relieving the hyper-stabilization of kinetochore microtubule attachments caused by phospho-blocking mutations in the Ndc80 complex (33). Importantly, many of these weakened microtubule attachment phenotypes were also observed upon specific impairment of the microtubule-binding activity of the Ska complex. These *in vivo* observations are consistent with the idea that Ska complex makes a direct contribution to load-bearing at the kinetochore-microtubule interface. However, the load-bearing capacity of the Ska complex has been unclear, leaving open the possibility that its role is primarily indirect, via recruitment of PP1 phosphatase (10). We show here for the first time that the Ska complex alone can bear load on microtubule ends, that it can enhance Ndc80 complex-based coupling, and that this enhancement requires the Ska complex to bind both microtubules and Ndc80 complex. These observations strongly support the model that the Ska complex strengthens kinetochore-microtubule attachments by forming a load-bearing bridge between the Ndc80 complex and the microtubule (Figure 5E).

Cell biological (8, 34), biochemical (16, 35), and evolutionary analyses (36) have suggested that Ska complex might be a functional analogue of the yeast Dam1 complex. However, while the Dam1 complex oligomerizes into microtubule-encircling rings that enhance its tip-coupling performance (37–39), the Ska complex does not appear to form such rings (6). Nevertheless, we find that the Ska complex, like the Dam1 complex, can form load-bearing tip attachments on its own and increase the strength and stability of Ndc80 complex-based couplers. Thus, our results lend further support to the hypothesis that the human Ska and yeast Dam1 complexes are functional analogues.

Our crosslinking mass spectrometry shows that the Ska complex interacts with the coiled-coil regions of the Ndc80 complex through the Ska3 C-terminus, but the overall architecture of their assembly at the kinetochore is unknown. Recently, the yeast Ndc80 complex was reported to bind two Dam1 complex rings and perturbations to this two-ring binding created mitotic attachment defects (40). Further structural studies will be needed to determine the assembly stoichiometry and how the Ska complex binds coiled-coil regions along the entire 55-nm-long Ndc80 complex (41). Revealing how this load-bearing unit, comprised of the Ndc80 and Ska complexes, tracks with and captures the forces generated by a depolymerizing microtubule tip is critical to understanding how kinetochores translate microtubule depolymerization into chromosome segregation.

Interestingly, the enhancement of Ndc80 complex-based tip-attachments upon addition of Ska complex occurred selectively; only when the Ndc80 complex-based attachments were relatively weak. We speculate that this effect might arise because Ska complex preferentially strengthens Ndc80 complex binding to a particular region on the microtubule tip, such as the most terminal tubulin subunits, and that Ndc80 complex-based couplers under weakened conditions rely primarily on bonds in this region. Alternatively, the Ska complex-dependent enhancement might be sterically blocked when Ndc80 complexes bind microtubules with high cooperativity (20, 25). While further studies will be required to understand the molecular basis for this selectivity, the effect could explain how Ska complex specifically prevents kinetochore detachments during episodes of poleward movement in prometaphase (32).

The Ndc80 complex is highly phosphorylated in early mitosis by Aurora B which greatly reduces its affinity for microtubules to promote the correction of erroneous kinetochore-microtubule connections (30). The Ska complex localizes with the Ndc80 complex early in mitosis, starting in prometaphase (5, 14). We found that the Ska complex strengthens microtubule attachments of Ndc80 complexes with all 9 Aurora B phosphorylation sites mutated to phosphomimetic residues. This observation suggests that the Ska complex may antagonize the weakening of attachments by Aurora B during early mitosis. Like the Ndc80 complex, the Ska complex is phosphorylated by Aurora B. Mimicking this phosphorylation *in vivo* was found to reduce Ska complex kinetochore localization, delaying cell division and destabilizing kinetochore attachments (42). Correcting erroneous kinetochore microtubule attachments may require Aurora B to coordinately phosphorylate both the Ska and Ndc80 complexes to completely break microtubule connections.

## MATERIALS AND METHODS

### Protein Expression and Purification

The human Ska complex was generated from the coexpression of a plasmid encoding GST-Ska3 and a dicistronic plasmid encoding Ska1 and Ska2 in *E. coli* BL21(DE3) Rosetta 2 cells (Stratagene). The Ska complex was purified using a GS4B GST affinity column followed by anion exchange and size exclusion chromatography (GE Lifesciences). All Ska complex constructs had the GST tag removed during purification. During optical tweezers assays, only Ska complex constructs bound to beads contained an N-terminal His-tag on Ska1, whereas Ska complexes added in solution did not contain a His-tag. See Supplemental Table 3 for a list of the Ska complex proteins used in each experiment. Human Ndc80 complex was expressed from two dicistronic plasmids and purified by nickel affinity and size exclusion chromatography, as described previously (20). See Figure S6 for SDS-PAGE gels of purified Ska and Ndc80 complex constructs. See SI Material and Methods for further details.

### TIRF Microscopy

Ska complex microtubule binding was assessed using a custom TIRF microscope as described previously (43). Taxol-stabilized microtubules labeled at 1% with Alexa 647 were prepared by polymerizing ~20 µM tubulin at 37 °C for 30 minutes in: BRB80 (80 mM PIPES pH 6.8, 1 mM MgCl_2_, 1 mM EGTA), 1 mM GTP, 6 mM MgCl_2_ and 3.8% DMSO. After polymerization, 10 µM Taxol was added and microtubules were pelleted at 130,000g for 10 min at 37 °C. The microtubule pellet was resuspended in warm BRB80 with 10 µM Taxol. Flow chambers were assembled from glass slides and PEGylated coverslips as described previously (43). Flow chambers were washed with water and then incubated for 5-10 minutes with “rigor” kinesin in the reaction buffer: BRB80, 8 mg/mL BSA (bovine serum albumin), 2 mM DTT, 40 mM glucose, 200 µg/mL glucose oxidase and 35 µg/mL catalase. Labeled microtubules were incubated in the flow chamber for 5 minutes. Unbound microtubules were washed away and then Ska complex^Ska3-GFP^ diluted in reaction buffer was introduced into the flow chamber. 488 nm channel and 647 nm channels were simultaneously imaged at 10 Hz for 100-200 seconds using an EM-CCD camera controlled with iXon software (iXon 887-BI; Andor Technology). For particles bound to the slide, Ska complex^Ska3-GFP^ and yeast Ndc80 complex Nuf2-GFP were non-specifically bound to the coverslip in the reaction buffer and then free complexes were washed away. Particles bound to the slide versus bound to microtubules were measured at two different laser powers. Single particle tracking and particle intensity analysis was performed using custom Labview (National Instruments) and Matlab (Mathworks) software. Mean lifetimes and associated errors of microtubule binding were determined by bootstrapping analysis. Kaplan-Meier analysis of survival probability curves was performed using Matlab.

### Optical Tweezers Bead Motility Assay

Bead motility assays were performed on a custom optical tweezers microscope as described previously (39). Streptavidin coated 0.44 µm polystyrene beads were functionalized with biotinylated penta-His antibodies (Qiagen) and stored in BRB80 containing: 8 mg/mL BSA and 8 mM DTT. To coat beads, 14.2 pM of the penta-His functionalized beads were incubated for 1 hour at 4 °C with either Ska complex with a penta-His on Ska1 or Ndc80 complex with a penta-His tag on the C-terminus of Spc24 at concentrations of: 1-100 nM for Ska complex and 0. 2-15 nM Ndc80 complex. The number of molecules per bead was estimated from the molar ratios of beads to His-tagged coupler. The concentration of His-tagged couplers was calculated using BCA assays. The concentration of beads was estimated from the manufacturer’s specifications of fraction of polymer by mass in the stock solution, the polystyrene density and the average bead diameter (Spherotech). At the concentrations tested, the beads were not saturated, based on the manufacturer’s stated biotin binding capacity per bead and previous work showing a linear trend labeling beads with increasing concentrations of a GFP construct (19). Microscopy flow chambers were assembled using double-sided tape, glass slides and #1.5 coverslips. Coverslips used for Ska complex coupling experiments were plasma-cleaned whereas Ndc80 complex coupling experiments used coverslips that were acid-cleaned and passivated with PEG as described previously (43). A solution of 1 mg/mL biotinylated BSA was introduced into the chamber and incubated for 15 minutes at room temperature, followed by a wash with BRB80; then 1 mg/mL avidin was added and incubated for 5 minutes before a final wash with BRB80. 0.1 – 0.2 mg/mL biotinylated GMPCPP stabilized microtubule seeds were flown into the chamber and incubated for 5 minutes at room temperature before free seeds were washed away with a blocking solution of BRB80 containing: 8 mg/mL BSA, 1 mM GTP and 1 mg/mL κ-casein. The experiment was initiated by flowing in a solution of ~ 1 pM of coated beads, 5-8 µM of bovine tubulin in BRB80 containing: 8 mg/mL BSA, 1 mM GTP, 200 µg/mL glucose oxidase, 35 µg/mL catalase, 30 mM glucose, and 1 mM DTT. For in-solution Ska complex experiments, free untagged Ska complex was added to the reaction mixture at final concentrations of 10, 25 or 50 nM. The flow chamber was sealed with nail polish before data collection.

Rupture force and lifetime experiments were performed at 26 °C on a previously described optical tweezers instrument (39). Data was collected using in-house developed Labview software. For rupture force experiments, coated beads were bound to dynamic microtubule tips. A force of ~1-2 pN was applied opposite the microtubule tip and a bead was monitored for 20-30 seconds to verify it was tracking with the microtubule tip. A force ramp was applied that increased the force by 0.25 pN/s until the bead ruptured from the microtubule tip. For constant force experiments, an Ndc80 complex coated bead was bound to a dynamic microtubule tip and a constant force clamp of ~2 pN was applied. Beads were tracked with the microtubule tip until a detachment event or the bead stuck to the coverslip. Force versus time traces were calculated using in-house routines for the Igor software package (WaveMetrics). Rupture forces, detachment events and switching events were manually determined by examining force and position versus time traces. Survival probability curves were constructed and medians were calculated from data sets containing beads that ruptured from microtubule ends, beads that did not hold the initial preload force of 1-2 pN and beads that exceeded the maximum force of the instrument. Median errors were determined by bootstrapping analysis. Microtubule switching and coupler detachment rates were calculated by dividing the number of events by the total time in the assembly or disassembly state. Kaplan-Meier analysis of survival probability curves was performed using Matlab.

### Crosslinking Mass Spectrometry

Crosslinking mass spectrometry was performed as previously described (40, 44). In brief, crosslinking reactions were carried out at room temperature for 15 or 30 minutes with BS3 or EDC, respectively, and quenched with NH_4_HCO_3_. Microtubule pellets with crosslinked proteins were resuspended, reduced and alkylated prior to overnight trypsin digestion at room temperature. Mass spectrometry was performed on a Q-Exactive HF (Thermo-Fisher Scientific) in data dependent mode and spectra converted into mzML using msconvert from ProteoWizard (45). Kojak was used to identify crosslinked peptides and q-values were assigned using Percolator (46, 47). Raw MS spectra and processed results are available at: http://proxl.yeastrc.org/proxl/viewProject.do?project_id=49 using the ProXL web application (48). See SI Material and Methods for complete details.

## ACKNOWLEDGEMENTS

We thank the members of the Davis and Asbury labs for their helpful discussions. We thank Dr. Prasad Jallepalli for the gift of the Ska complex plasmids. This work was supported by National Institute of Health grants: F32 GM120912 to L.A.H, R01 GM040506 to T.N.D., P41 GM103533 to M.J.M and R01 GM079373 to C.L.A., and The David and Lucile Packard Fellowship 2006-30521 to C.L.A.

## SUPPLEMENTAL INFORMATION

**Supplemental Figure 1.**
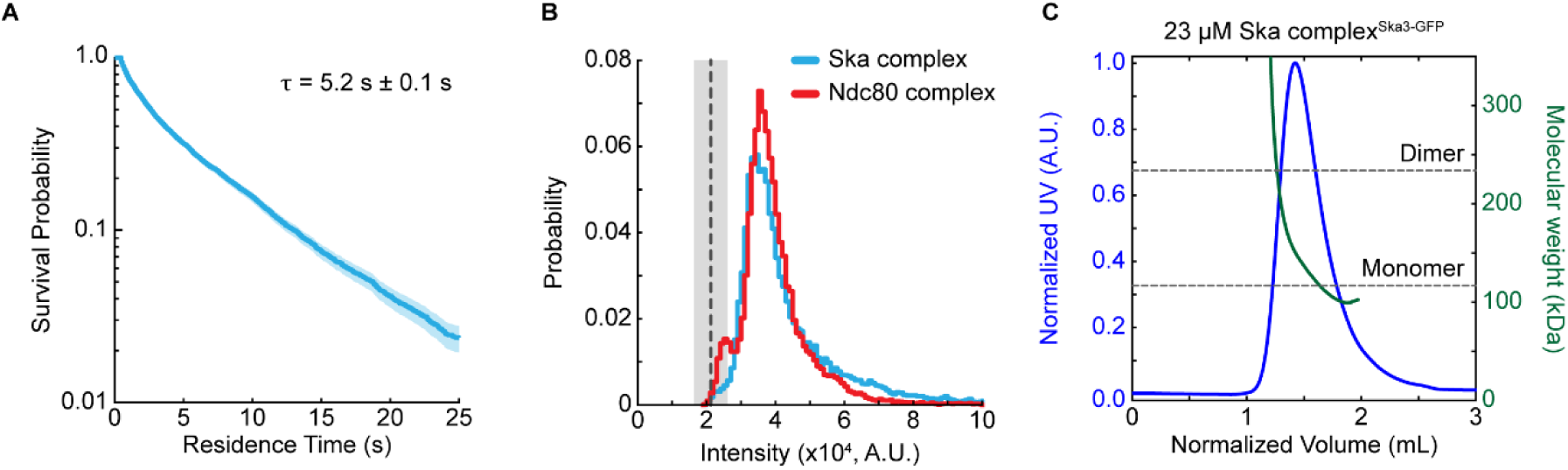
Ska complex^Ska3-GFP^ is monomeric at low concentrations and in a monomer-dimer equilibrium at micromolar concentrations. (A) Lifetime survival probability curve for 1 nM Ska complex^Ska3-GFP^ residence times on a microtubule lattice. Shaded region is 95% confidence interval from Kaplan-Meier analysis. Lifetime and error were calculated from bootstrap analysis. (B) Histogram of puncta intensities before photobleaching.125 pM Ska complex^Ska3-GFP^ (blue) or 125 pM yeast Ndc80 Complex^Nuf2-GFP^ (red) were added to coverslips and allowed to adhere before non-bound material was removed by washing. Particles were imaged by TIRF microscopy and their intensity measured. Black dashed line is the mean background intensity and shaded region shows the standard deviation of the background intensity after photobleaching. (C) Size exclusion chromatography multi-angle light scattering traces (SEC-MALS) of Ska complex^Ska3-GFP^ at 23 µM. Gray dashed lines indicated predicted molecular weights of the monomer and dimer.

**Supplemental Figure 2.**
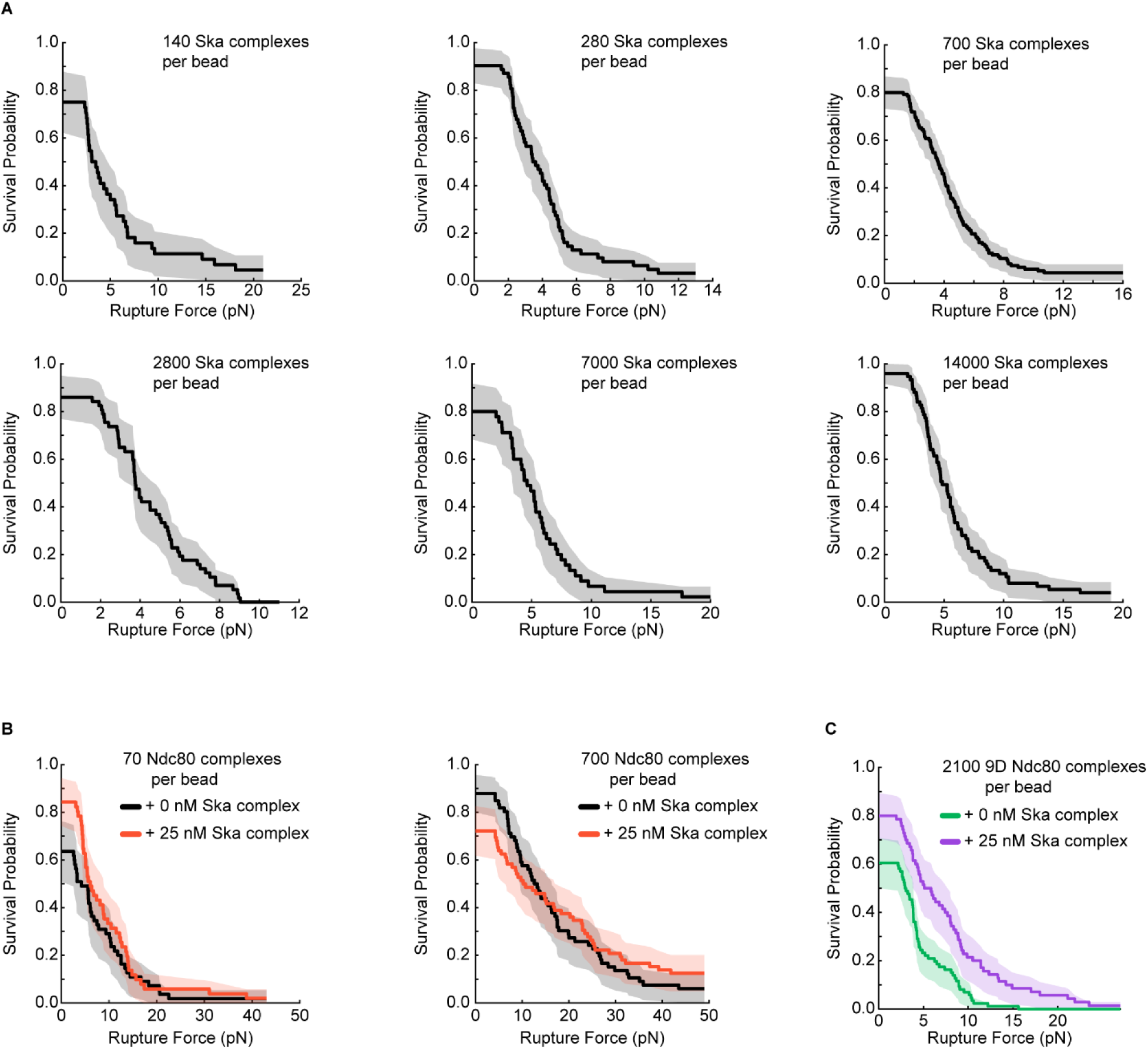
Rupture Force Survival Probability Plots. (A) Survival probability plots of rupture force data used to calculate the medians given in Figure 1G. (B) Survival probability plots of rupture force data used to calculate the medians given in Figure 3E with Ndc80 complex on the bead without (black) or with (orange) 25 nM Ska complex in solution. (C) Survival probability plots of rupture force data used to calculate the medians in Figure 3E with 9D Ndc80 complex on the bead without (green) or with (purple) 25 nM Ska complex in solution. All shaded regions on survival probability plots are 95% confidence intervals from Kaplan-Meier analysis.

**Supplemental Figure 3.**
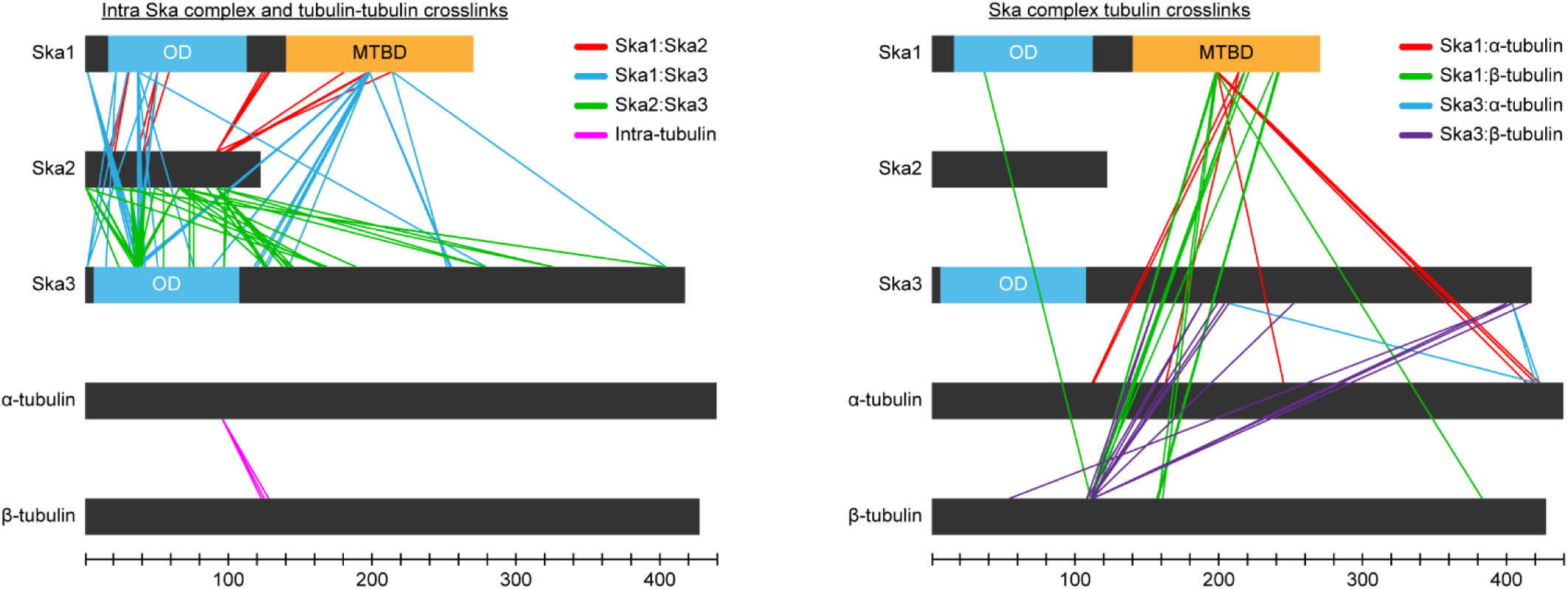
All crosslinks between Ska complex and microtubules. (Left) Crosslinks identified between Ska complex subunits and crosslinks identified between α- and β-tubulins. (Right) Crosslinks identified between Ska complex subunits and α- or β-tubulins. Crosslinking reaction with Ska complex and Taxol-stabilized microtubules was performed for 30 minutes with the amine to carboxyl crosslinker EDC. All data is available at http://proxl.yeastrc.org/proxl/viewProject.do?project_id=49

**Supplemental Figure 4.**
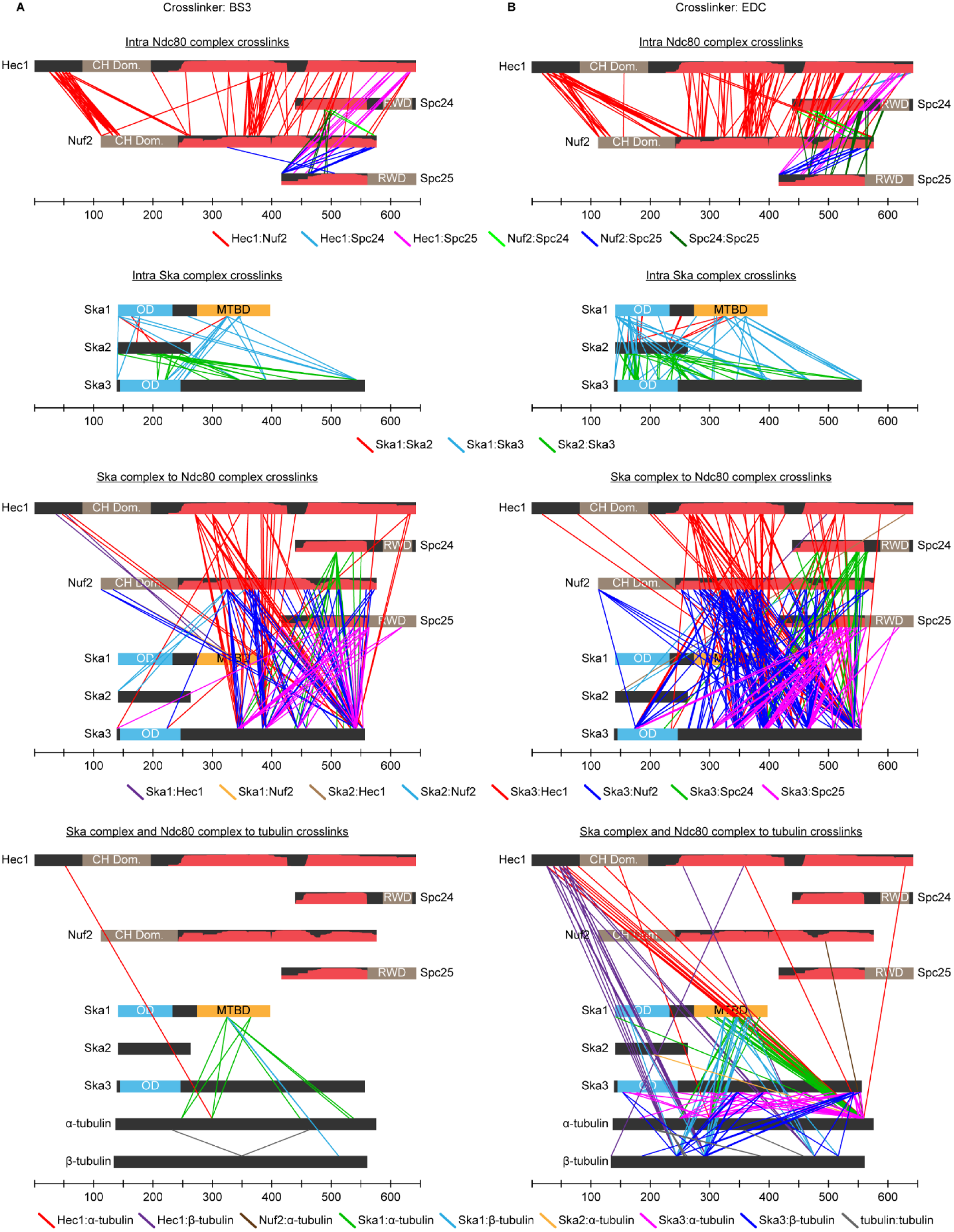
All crosslinks between Ska complex, Ndc80 complex and microtubules. (A) All crosslinks identified from the reaction with Ska complex, Ndc80 complex and Taxol-stabilized microtubules performed for 15 minutes with the amine to amine crosslinker BS3. (B) All crosslinks identified from the reaction with Ska complex, Ndc80 complex and Taxol-stabilized microtubules performed for 30 minutes with the amine to carboxyl crosslinker EDC. All data is available at http://proxl.yeastrc.org/proxl/viewProject.do?project_id=49

**Supplemental Figure 5.**
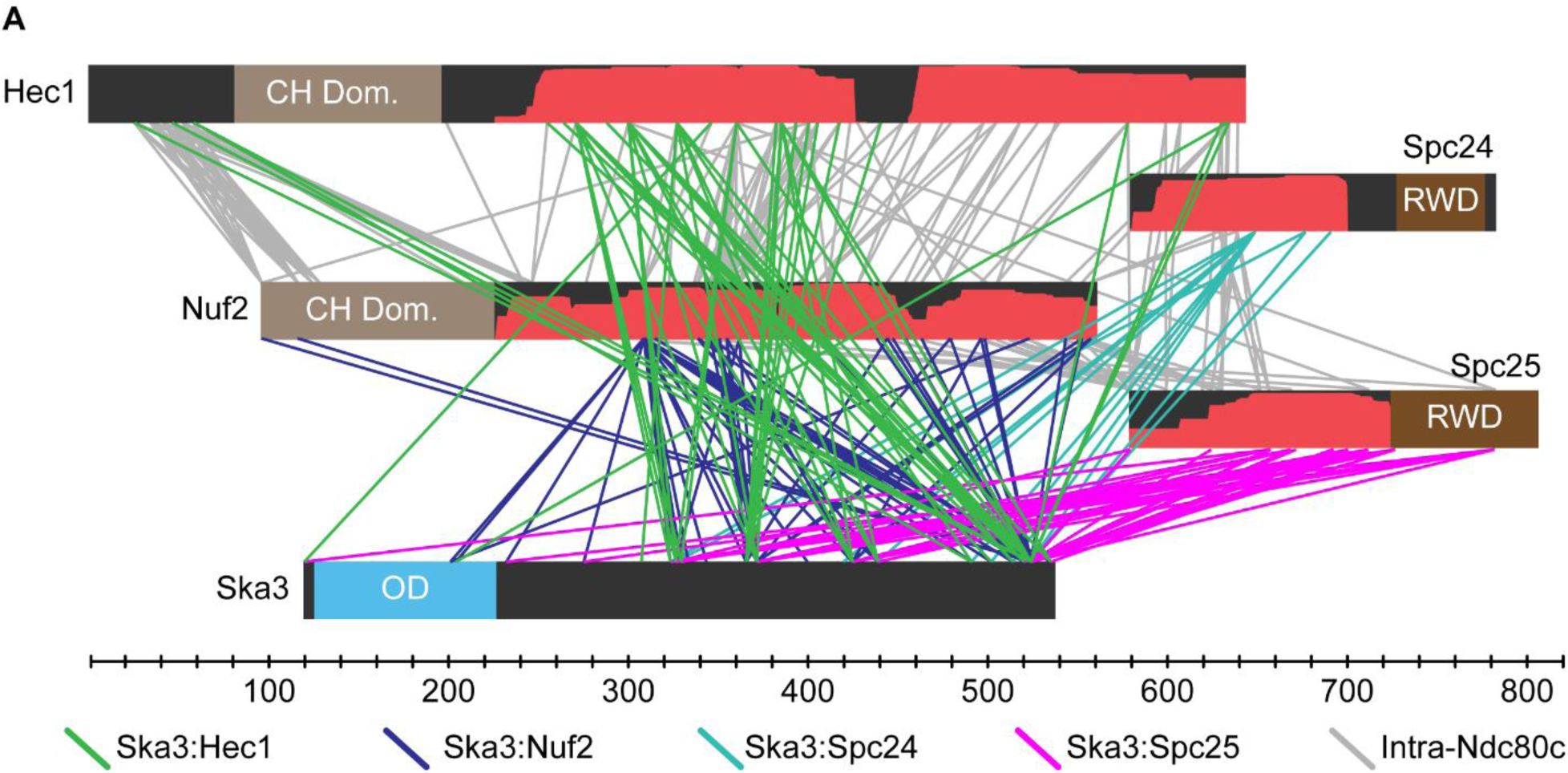
Ska complex^Ska1 ΔMTBD^ crosslinks with Ndc80 complex through the Ska3 C-terminus. (A) Crosslinks identified between Ska3 of Ska complex^Ska1 ΔMTBD^ and Ndc80 complex. Crosslinking reaction with Ska complex^Ska1 ΔMTBD^, Ndc80 complex and Taxol-stabilized microtubules was performed for 15 minutes with the amine to amine crosslinker BS3. Ska1, Ska2 and tubulin crosslinks are not shown for clarity; see http://proxl.yeastrc.org/proxl/viewProject.do?project_id=49 for all data.

**Supplemental Figure 6.**
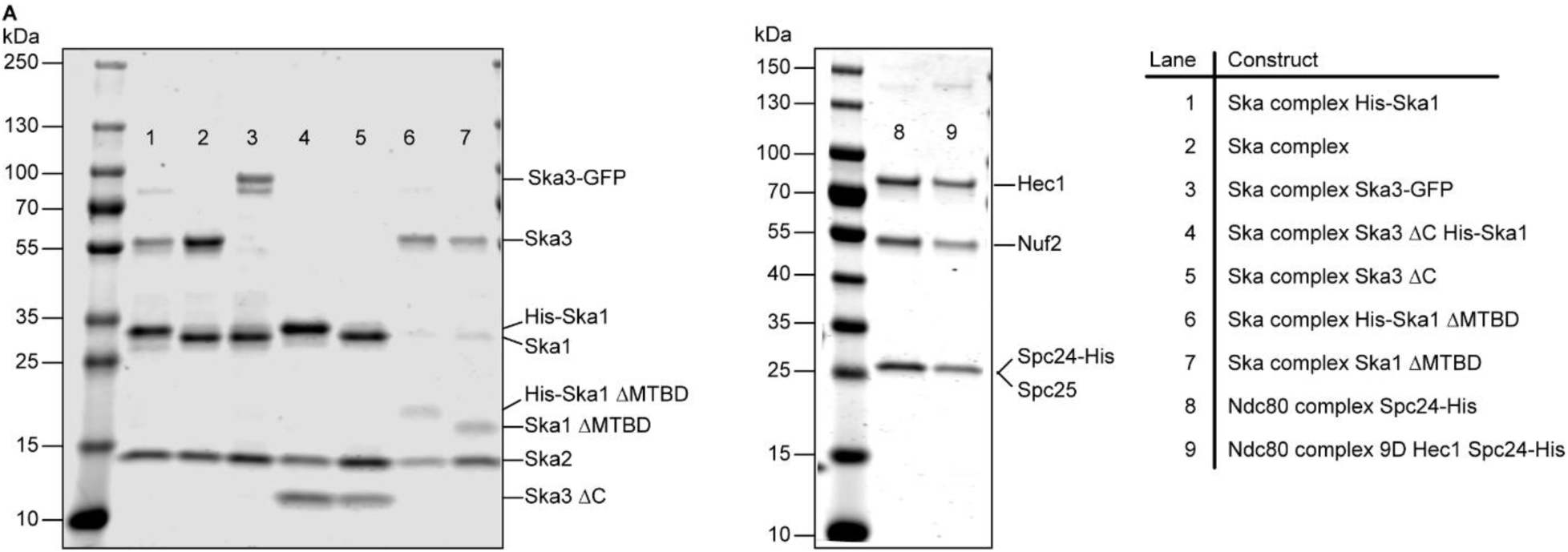
Purified recombinant Ska and Ndc80 complexes. (A) SDS-PAGE gels of purified recombinant Ska and Ndc80 complexes. Samples were run on 4-20% gradient polyacrylamide gels.

**TABLE S1.**

| Binding Elements | Median Rupture Force (pN) | Median Standard Error (pN) | N | # beads that reached max force | # Beads that did not hold preload |
| --- | --- | --- | --- | --- | --- |
| **Ska Complex Alone** |  |  |  |  |  |
| 140 Ska complexes per bead | 3.25 | 0.61 | 44 | 2 | 11 |
| 280 Ska complexes per bead | 3.52 | 0.38 | 62 | 2 | 6 |
| 700 Ska complexes per bead | 3.63 | 0.30 | 139 | 7 | 30 |
| 2800 Ska complexes per bead | 3.75 | 0.33 | 57 | 0 | 8 |
| 7000 Ska complexes per bead | 4.61 | 0.56 | 45 | 1 | 9 |
| 14100 Ska complexes per bead | 4.84 | 0.41 | 75 | 3 | 3 |
| 700 Ska complexes per bead + 10 nM Ska complex in solution | 3.38 | 0.34 | 110 | 0 | 22 |
| 700 Ska complexes Ska1 ΔMTBD per bead | ND | ND | 51 | 1 | ND |
| 700 Ska complexes Ska3 ΔC per bead | 2.74 | 0.24 | 88 | 0 | 25 |
| **Ndc80 Complex and Ska Complex** |  |  |  |  |  |
| 30 Ndc80 complexes per bead | 3.03 | 0.64 | 155 | 3 | 63 |
| 70 Ndc80 complexes per bead | 4.19 | 1.61 | 55 | 1 | 20 |
| 700 Ndc80 complexes per bead | 12.9 | 1.56 | 66 | 4 | 8 |
| 30 Ndc80 complexes per bead + 25 nM Ska complex in solution | 8.11 | 0.83 | 64 | 1 | 9 |
| 70 Ndc80 complexes per bead + 25 nM Ska complex in solution | 6.05 | 1.40 | 51 | 1 | 8 |
| 700 Ndc80 complexes per bead + 25 nM Ska complex in solution | 10.5 | 3.15 | 72 | 9 | 20 |
| 700 Ndc80 complexes 9D per bead | < 1 | 0.5 | 97 | 0 | 51 |
| 700 Ndc80 complexes 9D per bead + 25 nM Ska complex in solution | 5.61 | 0.51 | 75 | 1 | 21 |
| 2100 Ndc80 complexes 9D per bead | 3.00 | 0.51 | 86 | 0 | 34 |
| 2100 Ndc80 complexes 9D per bead + 25 nM Ska complex in solution | 5.47 | 1.12 | 70 | 1 | 14 |
| 700 Ndc80 complexes ΔN per bead | < 1 | ND | 74 | 0 | 53 |
| 700 Ndc80 complexes ΔN per bead + 25 nM Ska complex in solution | 5.88 | 1.22 | 59 | 1 | 18 |
| 30 Ndc80 complexes per bead + 25 nM Ska complex Ska3 ΔC in solution | 4.63 | 0.99 | 71 | 21 | 1 |
| 30 Ndc80 complexes per bead + 25 nM Ska complex Ska1 ΔMTBD in solution | 3.27 | 0.50 | 94 | 35 | 0 |
| ND = Not Determined |  |  |  |  |  |

**TABLE S2.**

|  |  |  |  |  |
| --- | --- | --- | --- | --- |
| **Catastrophe Rate** |  |  |  |  |
| **Binding Elements** | **Total Assembly Time (hr)** | **# Catastrophes** | **Catastrophe Rate (hr-1)** | **Catastrophe Rate Error (hr-1)** |
| Ndc80 complex | 3.77 | 10 | 2.65 | 0.84 |
| Ndc80 complex + Ska complex | 3.73 | 26 | 6.97 | 1.37 |
| **Rescue Rate** |  |  |  |  |
| **Binding Elements** | **Total Disassembly Time (hr)** | **# Rescues** | **Rescue Rate (hr-1)** | **Rescue Rate Error (hr-1)** |
| Ndc80 complex | 0.10 | 2 | 20.0 | 14.1 |
| Ndc80 complex + Ska complex | 0.17 | 16 | 94.1 | 23.5 |
| **Detachment Rate** |  |  |  |  |
| **Assembling Microtubules** |  |  |  |  |
| **Binding Elements** | **Total Assembly Time (hr)** | **# Detachments** | **Detachment Rate (hr-1)** | **Detachment Rate Error (hr-1)** |
| Ndc80 complex | 3.77 | 28 | 7.43 | 1.40 |
| Ndc80 complex + Ska complex | 3.73 | 25 | 6.70 | 1.34 |
| **Detachment Rate** |  |  |  |  |
| **Disassembling Microtubules** |  |  |  |  |
| **Binding Elements** | **Total Disassembly Time (hr)** | **# Detachments** | **Detachment Rate (hr-1)** | **Detachment Rate Error (hr-1)** |
| Ndc80 complex | 0.10 | 10 | 100 | 31.6 |
| Ndc80 complex + Ska complex | 0.17 | 9 | 52.9 | 17.6 |
| **Applied Force** |  |  |  |  |
| **Binding Elements** | **# of Beads Tested** | **# of Beads holding Preload** | **Mean Force (pN)** | **Force SEM (pN)** |
| Ndc80 complex | 192 | 53 | 1.82 | 0.05 |
| Ndc80 complex + Ska complex | 104 | 60 | 1.81 | 0.04 |
| Error = $\sqrt{N}/\text{time}$ | | | | |
| SEM = standard error of the mean |  |  |  |  |

**TABLE S3.**

| **Optical Tweezers Experiments** | **Ska Complex Proteins\*** | **Figures** |
| --- | --- | --- |
| Rupture Force with Ska complex on the bead | His-Ska1, Ska2, Ska3 | Figure 1, 2 and S2 |
| Rupture Force with Ska complex<sup>Ska3ΔC</sup> on the bead | His-Ska1, Ska2, Ska3ΔC | Figure 2 |
| Rupture Force with Ska complex<sup>Ska1ΔMTBD</sup> on the bead | His-Ska1ΔMTBD, Ska2, Ska3 | Figure 2 |
| Rupture Force with Ndc80 complex on the bead and Ska complex in solution | Ska1, Ska2, Ska3 | Figure 3, 5 and S2 |
| Force Clamping with Ndc80 complex on the bead and Ska complex in solution | Ska1, Ska2, Ska3 | Figure 4 |
| Rupture Force with Ndc80 complex on the bead and Ska complex<sup>Ska3ΔC</sup> in solution | Ska1, Ska2, Ska3ΔC | Figure 5 |
| Rupture Force with Ndc80 complex on the bead and Ska complex<sup>Ska1ΔMTBD</sup> in solution | Ska1ΔMTBD, Ska2, Ska3 | Figure 5 |
| **Crosslinking Experiments** |  |  |
| Ska complex with Taxol-microtubules | His-Ska1, Ska2, Ska3 | Figure 2 and S3 |
| Ndc80 complex with Ska complex and Taxol-microtubules | Ska1, Ska2, Ska3 | Figure 2 and S4 |
| Ndc80 complex with Ska complex<sup>Ska1ΔMTBD</sup> and Taxol-microtubules | Ska1ΔMTBD, Ska2, Ska3 | Figure S5 |
| **TIRF Microscopy Experiments** |  |  |
| Ska complex<sup>Ska3-GFP</sup> | Ska1, Ska2, Ska3-GFP | Figure 1 and S1 |
| **SEC-MALS Experiments** |  |  |
| Ska complex<sup>Ska3-GFP</sup> | Ska1, Ska2, Ska3-GFP | Figure S1 |
| * All Ska3 proteins were expressed as GST-Ska3 and the GST was removed with TEV cleavage during purification |  |  |

## SI MATERIALS AND METHODS

### Protein Expression and Purification

Human Ska complex was generated from a dicistronic pRSF plasmid encoding Ska1 and Ska2 and a pGEX plasmid encoding GST-Ska3 (a gift from Dr. Prasad Jallepalli). A Tobacco Etch Virus (TEV) protease site was introduced between the GST tag and Ska3 for affinity tag removal. His-tagged versions of the Ska complex were generated by introducing a penta-Histidine tag to the N-terminus of Ska1. The amino acid sequence for Ska complex mutant constructs are as follows: Ska complex^Ska3 ΔC^ is full length Ska1 and Ska2 with Ska3 residues 1-101; Ska complex^Ska1 ΔMTBD^ is full length Ska2 and Ska3 with Ska1 residues 1-131; Ska complex^Ska3-GFP^ is full length Ska1 and Ska2 with GFP attached to the C-terminus of full length Ska3. Human Ndc80 complex was generated from two dicistronic plasmids encoding Hec1 and Nuf2 or Spc24-His and Spc25. GFP was attached to the C-terminus of Nuf2 for S. *cerevisiae* Ndc80 complex GFP (19). Ndc80 complex 9D contains the following mutations: S4D, S5D, S8D, S15D, S44D, T49D, S55D, S62D, and S69D. Ndc80 complex ΔN is residues 81-642 of Hec1 and full length Nuf2, Spc24-His and Spc25. Plasmids and mutations were generated using standard molecular cloning procedures and Quickchange mutagenesis (Stratagene).

Ska1, Ska2 and Ska3 were coexpressed in BL21(DE3) Rosetta 2 *E. coli* cells (Stratagene) grown for 12-16 hours at 22 °C after induction with 0.3 mM IPTG. Cells were lysed using a French Press in 50 mM sodium phosphate buffer, pH 8.0, containing 300 mM NaCl, 2 mM DTT, 1 mM PMSF, 1 mM EDTA, 0.1% Tween-20, Benzonase nuclease and protease inhibitors. The lysate was clarified by a 40,000g spin for 20 minutes in a JA.25 rotor at 4 °C and loaded on a GS4B column (GE Life Sciences) at ~ 1 mL/min and 4°C. The GS4B column was washed for 3-5 column volumes with 50 mM sodium phosphate buffer, pH 8.0, containing 500 mM NaCl and 1 mM DTT before equilibration with a TEV cleavage buffer of 50 mM Tris buffer, pH 7.0, 150 mM NaCl, 1 mM EDTA, and 1 mM DTT. On-column TEV cleavage was performed 12-16 hours at 4 °C to cleave the GST from Ska3. The cleaved Ska complex was collected and concentrated to < 2mL using a 50,000 MW cutoff concentrator. Ska complex was further purified over a Superdex 200 16/60 column (GE Life Sciences) into a final buffer of 20 mM HEPES, pH 7.0, 150 mM NaCl, 5% glycerol and 1 mM DTT and then flash frozen in liquid nitrogen before storage at −80 °C. See Figure S6 for SDS-PAGE gels of each purified Ska complex construct.

Ndc80 complex was expressed and purified as described previously (20). In brief, Hec1, Nuf2, Spc24 and Spc25 were coexpressed in Rosetta 2 cells at 22 °C for 12-16 hours. After lysis and clarification, Ndc80 complex was purified using Ni-NTA affinity chromatography followed by size-exclusion chromatography using a Superdex 200 16/60 column. Ska and Ndc80 complex mutants were expressed and purified the same as wild-type. Protein concentrations were determined using a bicinchoninic acid assay. See figure S6 for SDS-PAGE gels of purified Ndc80 complex constructs.

### Crosslinking Mass Spectrometry

Crosslinking mass spectrometry was carried out as previously described (40, 44). For Ska complex on microtubules a 100 µL reaction in BRB80 was set up containing 40 µg of Ska complex plus 10 µg of Taxol-stabilized microtubules (made as described above). Reactions were incubated for 5 minutes at room temperature before adding 7.5 µL of 145 mM EDC and 3.75 µL of 145 mM Sulfo-NHS (dissolved in BRB80). Crosslinking was performed at room temperature for 30 minutes before quenching by addition of 5 µL 1M ammonium bicarbonate. The quenched reaction was centrifuged at 130,000g in a TLA100 rotor for 10 minutes at 37 °C and the resulting pellet was resuspended in 100 µL ice cold PBS containing 50 mM ammonium bicarbonate plus 1 µL 2M BME. The reaction was reduced for 30 minutes at 42 ^o^C with 10 mM DTT and alkylated for 30 minutes at room temperature with 15 mM iodoacetamide. Trypsin digestion was performed at room temperature overnight with shaking at a substrate to enzyme ratio of 60:1 prior to acidification with 5 M HCl. For Ska complex and Ska complex^Ska1 ΔMTBD^ plus Ndc80 complex on microtubules, 100 µL reactions in BRB80 containing 5 µg of Ska complex, 5 µg of Ndc80 complex and 5 µg of Taxol-stabilized microtubules were set up as before. EDC reactions contained 7.5 µL of 145 mM EDC and 3.75 µL of 145 mM Sulfo-NHS (dissolved in BRB80) and were crosslinked for 30 minutes at room temperature. BS3 reactions contained 3 µL 14.5 mM BS3 (dissolved in BRB80) and were crosslinked for 15 minutes at room temperature. Quenched reactions were centrifuged as before and pellets resuspended in 50 µL ice cold 20 mM ammonium bicarbonate with 0.5 µL 2M BME prior to reduction, alkylation and digestion as described above. Digested samples were acidified with 5 M HCl prior to being stored at −80 °C until analysis.

Mass spectrometry and data analysis was performed on a Q-Exactive HF (Thermo Fisher Scientific, Waltham, MA) as previously described (40, 44). Sample digest (0.8 - 1.5 µg) was loaded by autosampler onto a 150-µm Kasil fritted trap packed with Reprosil-Pur C18-AQ (3-µm bead diameter, Dr. Maisch) to a bed length of 2 cm. The trap was brought online with a 75-µm i.d. Pico-Frit column (New Objective) self-packed with 30 cm of Reprosil-Pur C18-AQ (3-µm bead diameter, Dr Maisch). Peptides were eluted from the column at 0.25 µL/min using a 120-minute acetonitrile gradient of 2% to 60%. Mass spectrometry was performed on a Q-Exactive HF (Thermo Fisher Scientific) in data dependent mode and spectra were converted into mzML using msconvert from ProteoWizard (45). Each sample was run 2-3 times and data were combined before analysis.

Crosslinked peptides were identified using Kojak version 1.4.3 (46) available at (http://www.kojak-ms.org). Percolator version 2.08 was used to assign a statistically meaningful q values to Kojak identifications (47). Target databases consisted of all proteins identified in the sample analyzed; decoy databases consisted of the corresponding set of reversed protein sequences. Data presented here were filtered to show hits to the target proteins that had a Percolator assigned peptide level q value ≤ 0.01. The complete, unfiltered list of all PSMs and their Percolator assigned q values, is available, along with the raw MS spectra and search parameters used, on the ProXL web application (48) at: http://proxl.yeastrc.org/proxl/viewProject.do?project_id=49

### Multi-Angle Light Scattering

Size exclusion chromatography coupled with multi-angle light scattering was performed on a Superdex 200 Increase 3.2/100 column (GE Life Sciences) in a buffer of 20 mM HEPES pH 7.0, 150 mM NaCl, 5% glycerol and 1 mM DTT. Elution from the size exclusion column was monitored by UV absorption at 280 nm, light scattering at 650 nm (miniDAWN Treos II, Wyatt Technologies) and differential refractometry (Optilab T-rEX, Wyatt Technologies). Data was analyzed using ASTRA software (Wyatt Technologies). Bovine Serum Albumin (BSA) at 5 mg/mL was used to calibrate the instrument specific parameters used during analysis.

